# Identification of Biochemical and Molecular Markers of Early Aging in Childhood Cancer Survivors

**DOI:** 10.1101/2021.04.01.438017

**Authors:** Silvia Ravera, Tiziana Vigliarolo, Silvia Bruno, Fabio Morandi, Danilo Marimpietri, Federica Sabatini, Monica Dagnino, Andrea Petretto, Martina Bartolucci, Monica Muraca, Eleonora Biasin, Riccardo Haupt, Marco Zecca, Franca Fagioli, Daniela Cilloni, Marina Podestà, Francesco Frassoni

**Affiliations:** Stem Cell Laboratory and Cell Therapy Center, IRCCS Istituto Giannina Gaslini, Genoa, Italy.; Department of Experimental Medicine, University of Genoa, Genoa, Italy.; Core Facilities-Clinical Proteomics and Metabolomics, IRCCS Istituto Giannina Gaslini, Genoa, Italy.; Epidemiology and Biostatistics Unit and DOPO Clinic, IRCCS Istituto Giannina Gaslini, Genoa, Italy.; Department of Pediatric Onco-Haematology, Regina Margherita Children’s Hospital, University of Turin - Turin, Italy; Pediatric Hematology Oncology, Fondazione IRCCS Policlinico San Matteo, Pavia, Italy.; Department of Clinical and Biological Sciences, School of Medicine, University of Turin, Turin, Italy.; Department of Mathematics (DIMA), University of Genoa, Genoa, Italy

## Abstract

**Purpose:** Survival rates of Childhood Cancer Patients have improved tremendously over the past four decades. However, cancer treatments are associated with an increased risk of developing an anticipated onset of chronic diseases typical of aging. Thus, we aimed to identify molecular/metabolic cellular alterations responsible for early aging in Childhood Cancer Survivors (CCS).

**Patients and Methods:** Biochemical, proteomic and molecular biology analyses were conducted on mononuclear cells (MNCs) isolated from peripheral blood of 196 CCS, comparing the results with those obtained on MNCs of 154 healthy subjects.

**Results:** Data demonstrate that CCS-MNCs show: i) inefficient oxidative phosphorylation associated with low energy status and a metabolic switch to lactate fermentation compared with age-matched normal controls; ii) increment of lipid peroxidation due to an unbalance among the oxidative stress production and the activation of the antioxidant defenses; (iii) significantly lower expression of genes and proteins involved in mitochondrial biogenesis and metabolism regulation, such as CLUH, PGC1-α, and SIRT6 in CCS, not observed in the age-matched healthy or elderly subjects. The application of a mathematical model based on biochemical parameters predicts that CCS have a biological age significantly increased by decades compared to the chronological age. Overall, the results show that the impact of chemo/chemoradiotherapy on mitochondria efficiency in 196 CCS was rather homogeneous, irrespective of cancer type, treatment protocols, and time elapsed from the end of the curative period.

**Conclusions:** Our study identifies some biochemical and molecular alterations possibly contributing to the pathophysiology of anticipated aging and metabolic deficiency described in CCS. These results may be useful in identifying approaches to restore the mitochondrial function, slowing down the aging and the associated pathological conditions in CCS.

## INTRODUCTION

Survival rates of Childhood Cancer Patients have improved tremendously over the past four decades since a 5-years survival rate is approaching over 80% ^1–3^. However, studies among Childhood Cancer Survivors (CCS) clearly show the risk of possible long term clinical complications related to chemotherapy, radiotherapy, or both ^4–6^. It is likely that the damage to normal tissues from cancer therapies diminishes physiological reserve, thus accelerating the process associated with aging, and increasing several accumulated stressors that impair the ability to restore physiologic turn-over and cellular homeostasis ^7^. In fact, several young adult CCS report frailty symptoms, consistent with early aging ^8–10^, as poor fitness, muscular weakness, and poor exercise tolerance ^11, 12^ as well as cognitive decline ^13, 14^, which appear decades earlier than expected ^4, 15–20^. The two most prevalent severe health conditions in these young cancer survivors are cardiovascular disease ^21^ and malignant neoplasms ^7^, conditions associated with aging in the general population. However, the cellular/molecular basis of all described symptoms and signs ^8, 9, 15^ remains missing. Aging is a process that involves several alterations such as telomere shortening, systemic inflammation, deregulated autophagy, oxidative stress, metabolic dysfunction, and epigenetic modifications ^22, 23^. In this scenario, senescent cells accumulate dysfunctional mitochondria that increase the reactive oxygen species (ROS) production, determining a negative effect on cellular bioenergetics ^24^, and driving cell senescence ^25^. The alteration of the oxidative phosphorylation (OxPhos) reduces the aerobic ATP production, inducing cells to switch to the anaerobic metabolism, probably to restore the cellular bioenergetics ^26, 27^.

In this contest, we generated a mathematical model able to predict the age of individuals with a mean absolute error of approximately 9 years ^28^. This “biological clock” is based on metabolic variables, which measure the modifications of glucose catabolism in the mononuclear cells (MNCs) of the healthy population (age between 5 and 106 years) ^28^. Using this model, we investigated whether CCS are suffering from a metabolic anticipated aging long term from the end of cancer therapies. In particular, we focused on mitochondria metabolism because their dysfunction determines the decrement of cellular energy availability and the imbalance between the generation of new mitochondria and their clearance, laying the foundation for aging.

## MATERIALS AND METHODS

### Reagents

All chemicals were purchased from Sigma Aldrich (St. Louis, MO, USA), unless otherwise indicated. Ultrapure water (Milli-Q; Millipore, Billerica, MA, USA) was used throughout. All other reagents were of analytical grade.

### Samples

All experiments were performed using MNCs isolated from peripheral blood (PB) with a standard gradient separation procedure, within 24 hours from collection. MNCs have been chosen because they are considered an excellent model to evaluate the health status of the entire organism^29^.

We studied 196 Childhood Cancer Survivors, after the end of treatments. This group of survivors had an age between 1-41 years old (y.o.): 103 with hematological malignancies (the commonest were ALL 60%, HD14% and AML 11.5%) and 93 with solid tumors (NB 59%, Wilms’ tumor14.7%, RSM 10.3%, Ewing sarcoma 3.8%, others 12.2%). All patients were given prolonged chemotherapy and/or radiotherapy according with clinical trials for pediatric malignancies effective at the onset of their diseases and were free of the disease at the time of the study. The median interval between completion of cancer treatment and blood sampling was 9 years (range: 2 months – 25 years). Age matched healthy donors with no cancer history (n. 79, age between 1-41 y.o.) as well as aged subjects (n. 75, age between 41 and >80 y.o.) were tested as controls (Table 1). This study was in accordance with the precepts established by the Helsinki Declaration and is approved by the Regional Ethics Committee of Liguria (N° 096REG2014) and all participants provided their written informed consent. For the participants under the age of 18 years, we have obtained the informed consent from a parent and/or legal guardian.

**Table 1.** Characteristics of CCS and control subjects.

| **Childhood Cancer Survivors** |  |  |  |  |  |  |
| --- | --- | --- | --- | --- | --- | --- |
| **Age** | **Solid Tumors** |  |  | **Haematological malignancies** |  |  |
| Years | Number of subjects | Female/Male | End of therapy (years) | Number of subjects | Female/Male | End of therapy (years) |
| <10<br>(Group 1) | 35 | 19/16 | 6 (1-9) | 32 | 25/7 | 2 (1-6) |
| 11-20<br>(Group 2) | 48 | 18/30 | 10 (3-17) | 38 | 16/22 | 4 (1-15) |
| 21-40<br>(Group 3) | 10 | 2/8 | 18 (4-25) | 33 | 21/12 | 9 (2-26) |
| **Total number** | **93** |  |  | **103** |  |  |
| **Controls** |  |  |  |  |  |  |
| **Age-matched** |  |  | **Aged controls** |  |  |  |
| Years | Number | Female/Male | Years | Number | Female/Male |  |
| <10 | 29 | 14/15 | 41-60 | 27 | 22/5 |  |
| 11-20 | 25 | 15/10 | 61-80 | 27 | 10/17 |  |
| 21-40 | 25 | 15/10 | >80 | 25 | 17/8 |  |
| **Total number** | **79** |  |  | **75** |  |  |

### Evaluation of oxygen consumption, ATP synthesis, P/O ratio, ATP/AMP ratio, and lactate dehydrogenase activity

In these experiments, MNCs samples were analysed within 24 hours from collection.

The oxygen consumption rate (OCR) was measured with an amperometric O_2_ electrode in a closed chamber, magnetically stirred, at 37 °C (Unisense, DK). For each assay, 200,000 cells were used. Sample was suspended in a medium containing: 137 mM NaCl, 5 mM KH_2_PO_4_, 5 mM KCl, 0.5 mM EDTA, 3 mM MgCl_2_ and 25 mM Tris–HCl, pH 7.4 and permeabilized with 0.03 mg/ml digitonin for 10 min. To stimulate the pathway composed by Complexes I, III and IV, 5 mM pyruvate plus 2.5 mM malate were added. To activate the pathway composed by Complexes II, III and IV 20 mM succinate was used ^30^.

ATP synthesis was measured by the highly sensitive luciferin/luciferase method. Assays was conducted at 37 °C, over 2 min, by measuring formed ATP from added ADP. 200,000 cells were added to the incubation medium (0.1 ml final volume) containing: 10 mM Tris-HCl pH 7.4, 50 mM KCl, 1 mM EGTA, 2 mM EDTA, 5 mM KH_2_PO_4_, 2 mM MgCl_2_, 0.6 mM ouabain, 0.040 mg/ml ampicillin, 0.2 mM p1p5-di(adenosine-5′) triphosphate, and the metabolic substrates: 5 mM pyruvate plus 2.5 mM malate or 20 mM succinate. The cells were equilibrated for 10 min at 37°C, then ATP synthesis has been induced by addition of 0.2 mM ADP. The ATP synthesis was measured using the luciferin/luciferase ATP bioluminescence assay kit CLSII (Roche, Basel, Switzerland), on a Luminometer (GloMax® 20/20 Luminometer – Promega, Wisconsin, USA). ATP standard solutions (Roche, Basel, Switzerland) in the concentration range 10^-10^ - 10^-7^ M was used for calibration ^28^.

Dividing the nmol of ATP produced by the nmol of oxygen consumed in one minute, we obtained the P/O ratio, which indicates the efficiency of the OxPhos metabolism. In coupled condition, when the oxygen consumption is correctly associated with the ATP synthesis, this value is around 2.5 or 1.5 in the presence of pyruvate + malate or succinate, respectively ^31^. Conversely, in case of uncoupled status, this value decreases proportionally to the grade of the OxPhos inefficiency.

The ATP and AMP quantification was based on the enzyme coupling method ^32^. 20 µg of total protein were used for both assays. Briefly, ATP was assayed following NADP reduction, at 340 nm. The medium contained: 100 mM Tris-HCl pH 8.0, 1 mM NADP, 10 mM MgCl_2_, and 5 mM glucose in 1 ml final volume. Samples were analysed spectrophotometrically before and after the addition of 4 µg of purified hexokinase plus glucose-6-phosphate dehydrogenase. AMP was assayed following the NADH oxidation at 340 nm. The medium contained: 100 mM Tris-HCl pH 8.0, 75 mM KCl, 5 mM MgCl_2_, 0.2 mM ATP, 0.5 mM phosphoenolpyruvate, 0.2 mM NADH, 10 IU adenylate kinase, 25 IU pyruvate kinase, and 15 IU of lactate dehydrogenase.

To assay the lactate fermentation flux, the activity of lactate dehydrogenase (LDH; EC 1.1.1.27) was measured at 25 °C on 20 µg of MNC homogenate. The reaction mixtures contained: 100 mM Tris-HCl pH 7.4, 0.2 mM NADH and 5 mM pyruvate^33^. NADH molar extinction coefficient was considered 0.622 mM^-1^ cm^-1^, at 340 nm. Enzymatic activity was expressed as mU/mg of total protein (nmol/min/mg of protein).

### ROS production and lipid peroxidation evaluation

To evaluate the ROS level, freshly isolated MNCs were washed and re-suspended in PBS and stained for 10 min at 37°C with 2’,7’-dichlorodihydrofluorescein diacetate (H2DCFDA) at a concentration of 5 µM (Thermo Fisher Scientific, Waltham, MA, USA). H2DCFDA is a non-fluorescent dye which is cleaved inside cells to 2’,7’-dichlorofluorescein (H2DCF). In the presence of oxidants, H2DFC is converted in turn to the fluorescent compound DCF. Samples were measured on a FacsCalibur flow cytometer (Becton Dickinson, San José, CA). The analysis was confined to viable cells only, after gating based on forward- and side-scatter characteristics. Ten thousand cells per sample were analyzed ^34^.

Malondialdehyde (MDA) concentration was evaluated as a marker of lipid peroxidation, by the thiobarbituric acid reactive substances (TBARS) assay ^35^. The TBARS solution contains: 15% trichloroacetic acid in 0.25 N HCl and 26 mM thiobarbituric acid. To evaluate the basal concentration of MDA, 600 μl of TBARS solution was added to 50 μg of total protein dissolved in l of milliQ water. The mix was incubated for 40 min at 95 °C. Afterward, the sample was μ centrifuged at 14000 rpm for 2 min and the supernatant was analysed spectrophotometrically, at 532 nm.

### Mitochondrial trans membrane potential by flow cytometry and confocal microscopy

The mitochondrial transmembrane potential of fresh isolated MNCs was analyzed both by flow cytometry, for evaluating the mean mitochondrial fluorescence intensity of the sample population, and by confocal microscopy for evaluating the intracellular mitochondrial distribution.

For flow cytometric evaluation, the fresh isolated MNCs were washed once with RPMI medium, incubated with 200 nM tetramethylrhodamine methyl ester (TMRM) (Invitrogen, Milan, Italy) for 10 minutes at 37°C and immediately measured on a FacsCalibur flow cytometer (Becton Dickinson, San José, CA). To exclude an unspecific staining, the same experiments has been conducted in the presence of 50 nM of Carbonyl cyanide-4-(trifluoromethoxy)phenylhydrazone (FCCP), an uncoupling molecule. The analysis was confined to viable cells only, after gating procedures based on forward- and side-scatter features. Ten thousand cells per sample were analyzed.

For fluorescence imaging, freshly isolated MNCs were incubated with 100 nM MitoTracker Deep Red-633 (ThermoFisher Scientific Inc.) for 15 min at 37 °C, placed on a coverslip and immediately analyzed using a Leica TCS SP2-AOBS confocal microscope (Leica Microsystem, Heidelberg, Germany) equipped with a 633 He-Ne laser and Leica specific software.

### Proteomic Setup

Samples were solubilized in 40 ul 2% SDC, 40 mM Chloroacetamide, 10 mM TCEP and 100 mM Tris-HCl pH 8 at 95°C for 10 min and sonicated with a Ultrasonic Processor UP200St (Hielscher), 3 cycles of 30 sec. Lysates samples were digested with 0.7 ug Trypsin and 0.3 μg LysC, overnight at 37 °C. Digested samples were processed by in-Stage Tip method using enclosed Stage Tip, containing 2 poly (styrene-divinylbenzene) reverse-phase sulfonate discs (SDB-RPS) ^36^. Elution was performed on an Ultimate 3000 RSLC with an EASY spray column (75 μm x 50 cm, 2 μm particle size, Thermo Scientific) at a flow rate of 250 nl/min with a 180 min non-linear gradient of 6-45% solution B (80% ACN, 20% H2O, 5% DMSO and 0.1% FA). Eluting peptides were analyzed using an OrbitrapVelos Pro mass spectrometer (Thermo Scientific Instruments) in positive ionization mode. Single MS survey scans were performed in the Orbitrap, recording a mass window between 375 and 1500 m/z using a maximal ion injection time of 50 ms. The resolution was set to 100,000 and the automatic gain control was set to 1,000,000 ions. The experiments were done in data-dependent acquisition mode with alternating MS and MS/MS experiments. A maximum of 10 MS/MS experiments were triggered per MS scan.

Data processing was performed by MaxQuant software version 1.6.1.0 ^37^. A false discovery rate was set at 0.01 for the identification of proteins and a minimum of 6 amino acids was required for peptide identification. Andromeda engine was used to search MS/MS spectra against Uniprot human database (release UP000005640_9606 August 2017).The intensity values were extracted and statistically evaluated using the ProteinGroup Table and Perseus software version 1.6.1.3^38^. Algorithm MaxLFQ was chosen for the protein quantification with the activated option ‘match between runs’ to reduce the number of the missing proteins.

The mass spectrometry proteomics data have been deposited to the ProteomeXchange Consortium via the PRIDE ^39^ partner repository with the dataset identifier PXD021960 (Reviewer account details: Username: Password: tFCOHhEx).

### qPCR analyses

RNA extraction from MNC cells from CCS or healthy donor was performed using the RNeasy Mini Kit (Qiagen, Milan, Italy) and quantified using a NanoDrop spectrophotometer (Nanodrop Technologies, Wilmington, DE). The cDNA was synthesized by using iScript cDNA Synthesis Kit (Bio-Rad Laboratories, Milan, Italy) starting from 1 μ were designed through Beacon Designer 2.0 Software (Bio-Rad Laboratories). CLUH primers sequences were TACATCATGGGCGACTACGC (forward primer) and GGCCAGGTGCATGTATTCCT (reverse primer); PGC1-alpha primers sequences were: CTGTGTCACCACCCAAATCCTTAT (forward primer) and TGTTCGAGAAAAGGACCTTGA (reverse primer); Sirt6 human primers sequences were: CCTCCTCCGCTTCCTGGTC (forward primer) and GTCTTACACTTGGCACATTCTTCC (reverse primer). Quantitative real-time PCR (qPCR) was performed in an iQ5 real-time PCR detection system (Bio-Rad Laboratories) using 2× iQ Custom Sybr Green Supermix (Bio-Rad Laboratories). Values were normalized on mRNA expression of human β-actin and HPRT (reference genes). Statistical analysis of the qPCR was performed using the iQ5 Optical System Software version 1.0 (Bio-Rad Laboratories) based on the 2− ΔΔCt method. The dissociation curve for each amplification was analyzed to confirm absence of unspecific PCR products.

### Western blot analysis

Expression of CLUH, PGC-1α and SIRT6 proteins was determined by Western blot (WB), using standard procedures. MNC were lysed using lysis Buffer (150 mM NaCl, 20 mM TRIS-HCl pH 7.4, 2 mM EDTA and 1% NP40) containing a protease inhibitor cocktail for mammalian cells (Sigma) and total protein was measured by Bradford assay ^40^. After SDS-PAGE, performed according to the standard method on 4-20% polyacrylamide gels, proteins were transferred to a nitrocellulose membrane (BioRad Laboratories). The membrane was blocked for 1 h with PBS- 0.1% Tween 20 (PBSt) containing 5% non-fat dry milk and incubated over night at 4°C with the following primary antibodies: anti-CLUH (1:1000, cod: A301-764A, Bethyl Lab), anti-PGC-1α (1:1000, cod: ab77210, Abcam), and anti-SIRT6 (1:1000, cod: ab88494, Abcam).

After washing with PBSt, the membrane was incubated with an anti-rabbit or anti-mouse IgG antibody conjugated with horse radish peroxidase (HRP) (BioRad Laboratories) and developed with Clarity Western ECL Substrate (BioRad Laboratories). Bands were detected and analyzed for density using an enhanced chemiluminescence system (Alliance 6.7 WL 20 M, UVITEC, Cambridge, UK) and UV1D software (UVITEC). Bands of interest were normalized for actin level in the same membrane.

### Statistical analysis

Experimental values are expressed as mean ± standard deviation. Comparison between different groups was performed by analysis of variance (ANOVA) for multiple comparisons followed by Bonferroni post hoc test; Student t test for paired or unpaired data, were used as appropriate. Analyses were performed using the GraphPad Prism version 5.00 statistical software (GraphPad Software Inc., La Jolla, CA, USA). Values of p< 0.05 were considered significant.

## RESULTS

### CCS-MNCs show an altered glucose metabolism and an increased oxidative stress production in comparison to the age-matched control cells

To assess the differences in mitochondrial metabolism between CCS and healthy control samples, we have evaluated several biochemical markers on MNCs isolated from peripheral blood. CCS have been stratified in three different groups, based on the age at analysis (Group 1=<10 y.o., Group = between 11 and 20 y.o., Group 3= between 21 and 40 y.o.) and compared with age-matched healthy subjects as well as adult and elderly controls (41-60, 61-80 and >80 y.o.) (Table 1).

The efficiency of oxidative phosphorylation (OxPhos) was evaluated in terms of P/O value, calculated as the ratio between the nmol of produced ATP and the nmol of consumed oxygen. This study was conducted in the presence of pyruvate + malate (Pyr+Mal), or succinate, to stimulate the pathways formed by complexes I, III, and IV and complexes II, III, and IV, respectively. As shown in Figure 1, Panel A, and Table 1 Supplementary, the P/O ratio after Pyr+Mal stimulation was lower in CCS patients affected either by solid or hematological tumors compared with the respective age-matched controls. In addition, the CCS P/O ratio resulted lower compared to the healthy population up to the group 41-60 y.o., while it was superimposable to subjects of 60 years older. Similar results were obtained evaluating the P/O ratio after succinate stimulation (Figure 1, Panel B, and Table 1 Supplementary).

**Figure 1.**
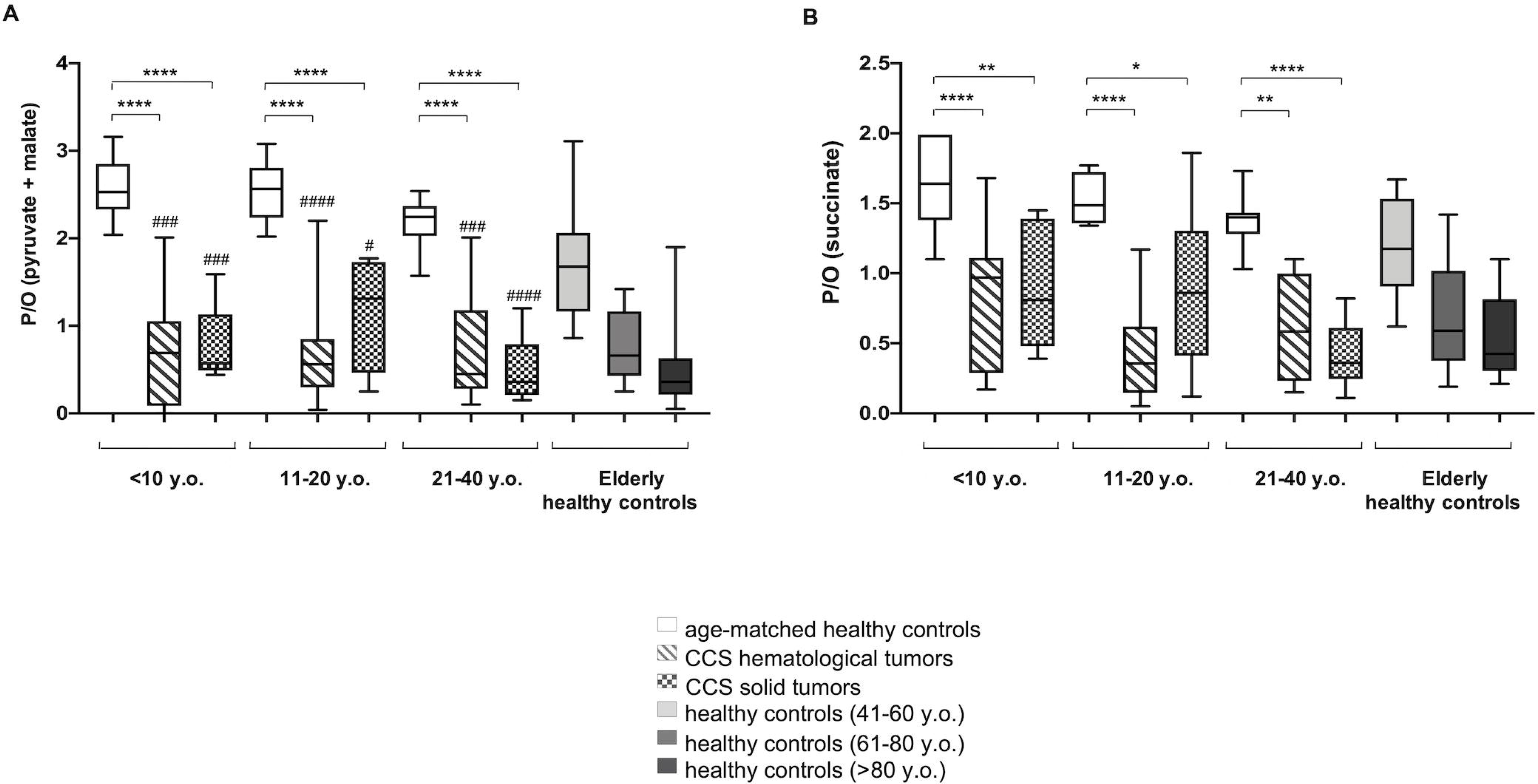
Evaluation of mitochondrial efficiency in MNC isolated from CCS, age-matched and elderly healthy controls. Panel A and B reports the P/O value, obtained as the ratio between ATP synthesis and oxygen consumption, in the presence of pyruvate + malate or succinate, respectively, in MNC isolated from: age-matched healthy control (<10 y.o. n= 17; 11-20 y.o. n=18; 21-40 y.o n=24), CCS hematological tumors (<10 y.o. n= 26; 11-20 y.o. n=53; 21-40 y.o n=10), CCS solid tumors (<10 y.o. n= 22; 11-20 y.o. n=19; 21-40 y.o n=14),adult healthy control (n= 21; 41-60 y.o.), and elderly healthy control (61-80 y.o. n=22 and>80 y.o. n=25). *, **, ***, and **** indicate a significant difference for p<0.05, 0.01, 0.001, and 0.0001, respectively, between CCS samples and age-matched healthy controls. #, ### and ####indicate a significant difference for p<0.05, 0.001, and 0.0001, respectively, between CCS samples and adult healthy controls (41-60 y.o.). No significant differences have been observed between CCS and elderly control (61-80, and >80 y.o.).

These data are confirmed by flow cytometric and confocal microscopy analysis of the mitochondrial membrane potential (MMP), evaluated with specific fluorescent probe TMRM. Data show a decrement of about 44% of MMP levels in CCS with respect to those observed in the healthy age-matched subjects (Figure 2, Panel A). The intracellular distribution of mitochondria in CCS cells appears to be altered as well. Although the pattern of mitochondria is not easily resolved in these cells that display a low cytoplasm/nucleus ratio, we observed a speckled distribution in CCS cells much different from the more uniformly distributed fluorescence in normal MNCs (Figure 2, Panel B), suggesting a loss in the mitochondrial network organization.

**Figure 2.**
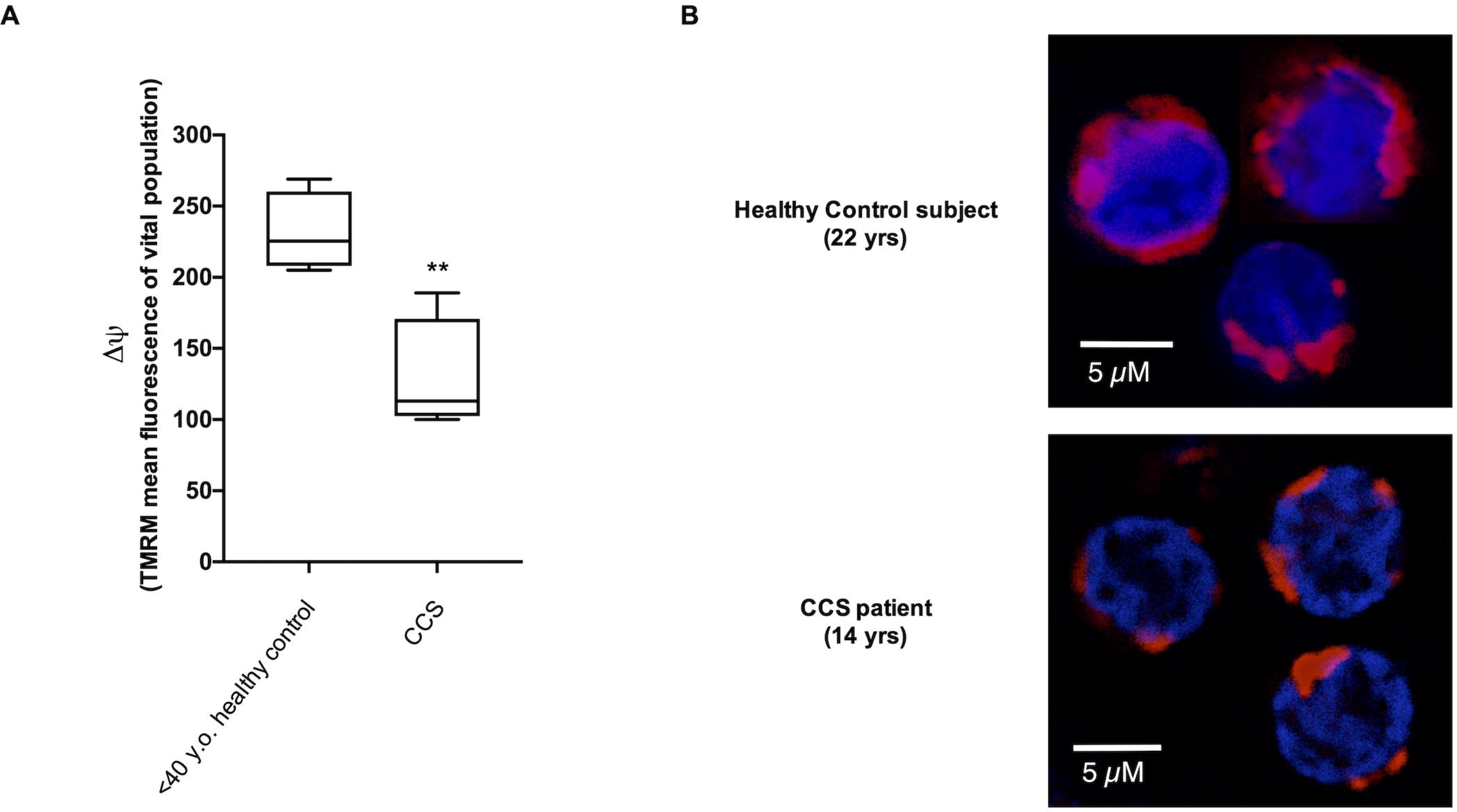
Evaluation of mitochondrial membrane potential in MNC isolated from CCS and age-matched healthy control. Panel A reports data on the mitochondrial transmembrane potential ΔΨ of MNC isolated from age-matched healthy controls and CCS patients, stained with the fluorescent probe TMRM and analyzed by flow cytometry. Data derive from five independent experiments and ** indicates p<0.001 (t test?). Panel B shows a representative confocal microscopy field of MNC isolated from one healthy subject (22 yrs) and from one CCS patient (14 yrs), stained with Mitotracker deep red 633 (virtual red color) as a marker of the mitochondrial membrane potential, and DAPI (blue color) to highlight the nuclei. Unlike in normal MNC, the mitochondria of CCS samples are distributed as single scattered spots.

An inefficient mitochondria metabolism is often associated with the increment of oxidative stress production ^41–43^. In CCS-MNC, the ROS production was ten-fold higher than in age-matched control cells (30.32 ± 5.39 vs 3.67 ± 0.90 mean fluorescence, respectively, p<0.001) (data not shown). Moreover, CCS patients, affected either by solid or hematological tumors, displayed levels of malondialdehyde (MDA), a marker of lipid peroxidation ^28^, higher than age-matched healthy controls, but similar to those reported for adult or elderly healthy control (Figure 3, Panel A, and Table 1 Supplementary).

**Figure 3.**
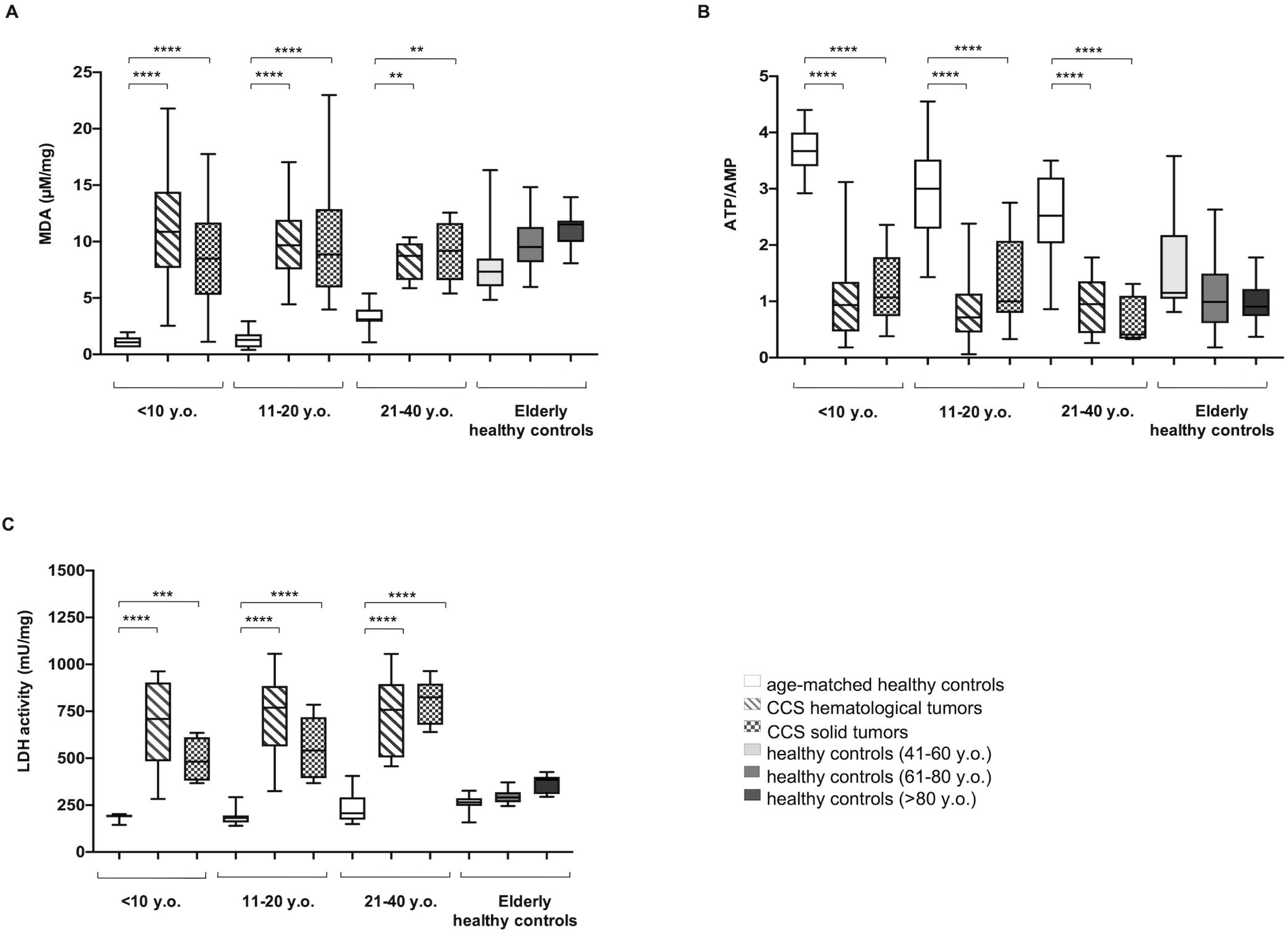
Lipid peroxidation, energy status and lactate dehydrogenase activity in MNC isolated from CCS, age-matched and elderly healthy controls. All data reported in this figure have been obtained using MNC isolated from: age-matched healthy control (<10 y.o. n= 17; 11-20 y.o. n=18; 21-40 y.o n=24), CCS hematological tumors (<10 y.o. n= 26; 11-20 y.o. n=53; 21-40 y.o n=10), CCS solid tumors (<10 y.o. n= 22; 11-20 y.o. n=19; 21-40 y.o n=14), adult healthy control (41-60 y.o n= 21;.), and elderly healthy control (61-80 y.o. n=22 and >80 y.o. n=25). Panel A shows the cellular level of malondialdehyde (MDA), a marker of lipid peroxidation. Data are expressed as μM/mg of total protein. Panel B reports the cellular energy status (ATP/AMP), evaluated as the ratio between the intracellular levels of ATP and AMP. Panel C shows the activity of lactate dehydrogenase (LDH), the marker of lactate fermentation. Data are expressed as mU/mg of total protein. **, ***, and **** indicate a significant difference for p<0.01, 0.001, and 0.0001, respectively, between CCS samples and age-matched healthy controls. No significant differences have been observed between CCS and adult and elderly controls (41-60, 61-80, and >80 y.o.).

Since OxPhos is the principal source of ATP production, we have evaluated the energy status of MNCs, analyzing the ATP/AMP ratio (Figure 3, Panel B, and Table 1 Supplementary). CCS showed a decreased energy status compared to age-matched healthy controls; such decrease was statistically significant at the young age-matched groups, while no significant difference has been observed between CCS samples and elderly healthy control.

We have previously demonstrated that mitochondria lose part of their energetic efficiency during aging, with a corresponding increase of lactate dehydrogenase (LDH), a marker of lactate fermentation, determining a metabolic switch from aerobic to anaerobic energy production ^28^. Here, we observed a significant increase of LDH activity in CCS-MNC either with hematological or solid tumors, in subjects of all groups we have tested (Figure 3, panel C and Table 1 Supplementary).

### The alteration of glucose metabolism is independent of the type of cancer and the respective therapy and does not ameliorate with time

The impact of cancer-specific treatments on glucose metabolism was investigated comparing the biochemical parameters of MNC from CCS who had been affected by different tumors (hematological vs solid): no significant differences were observed based on the type of cancer and the associated therapy. The efficiency of oxidative phosphorylation and the associated accumulation of lipid peroxidation as well as the alteration of the energy status and the activation of anaerobic glycolysis appeared similar in each sample from the 196 CCS (Figure 4).

In addition, our data show that there is no improvement in metabolic parameters over time since the last chemo/chemo-radiotherapy (range: 2 months – 25 years). In other words, it seems that the damage, once acquired, remains permanently (Figure 5).

**Figure 4.**
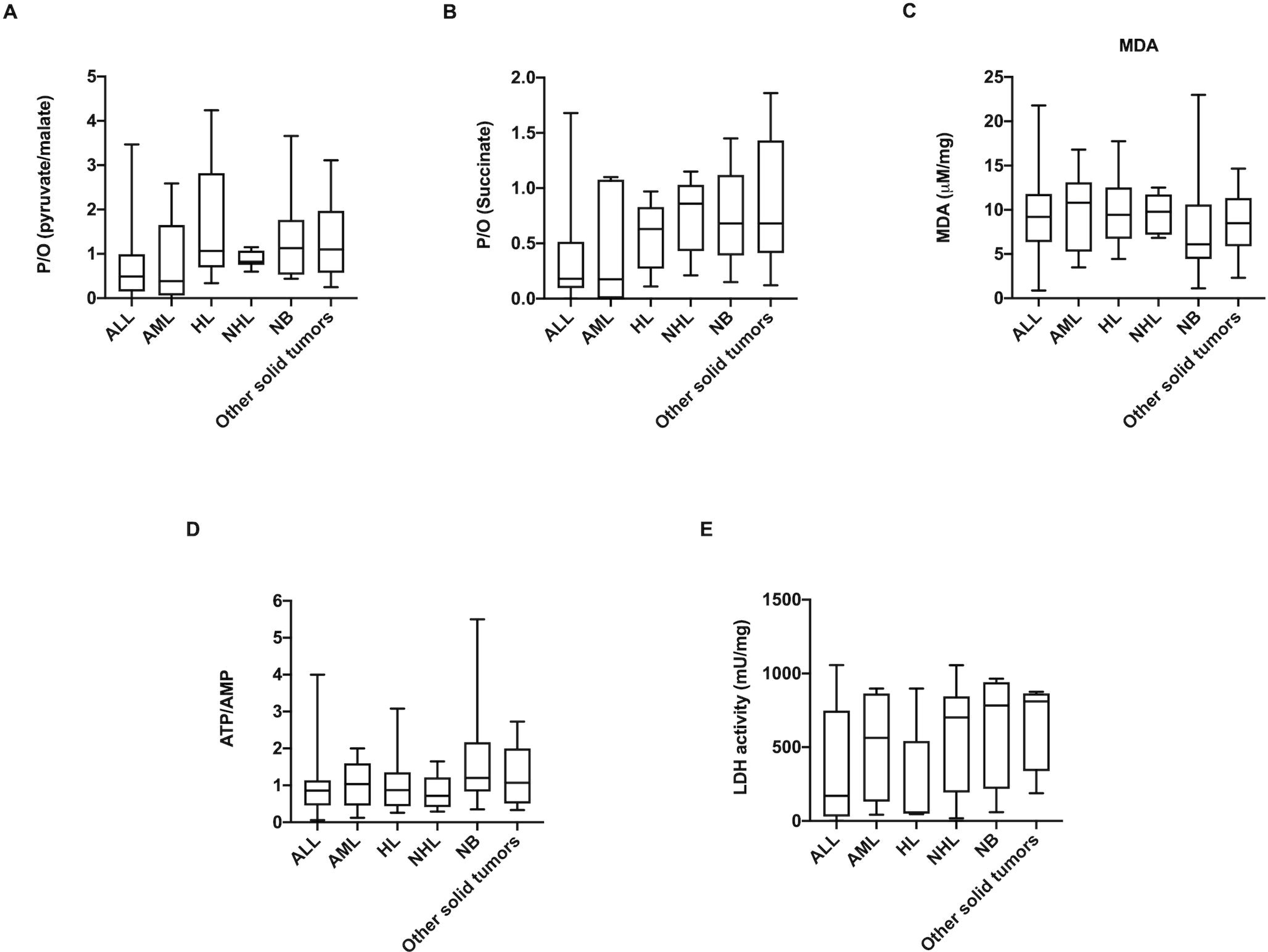
Comparison of glucose metabolism alteration in different type of tumors. All data reported in this figure were obtained in MNCs isolated from patients who survived the following types of tumors: acute lymphocytic leukemia (ALL; n=42), Acute myeloid leukemia (AML; n=15), Hodgkin lymphoma (HL; n=19), Non-Hodgkin lymphoma (NHL; n=13), Neuroblastoma (NB; n=34), and other solid tumors (n=21). Panel A and B reports the P/O value, obtained as the ratio between ATP synthesis and oxygen consumption, in the presence of pyruvate + malate or succinate, respectively. Panel C shows the MDA level, as marker of lipid peroxidation. Panel D reports the cellular energy status, expressed as the ATP/AMP ratio. Panel E shows the LDH activity, as marker of anaerobic glycolysis. No significant differences have been observed among the analyzed samples.

**Figure 5.**
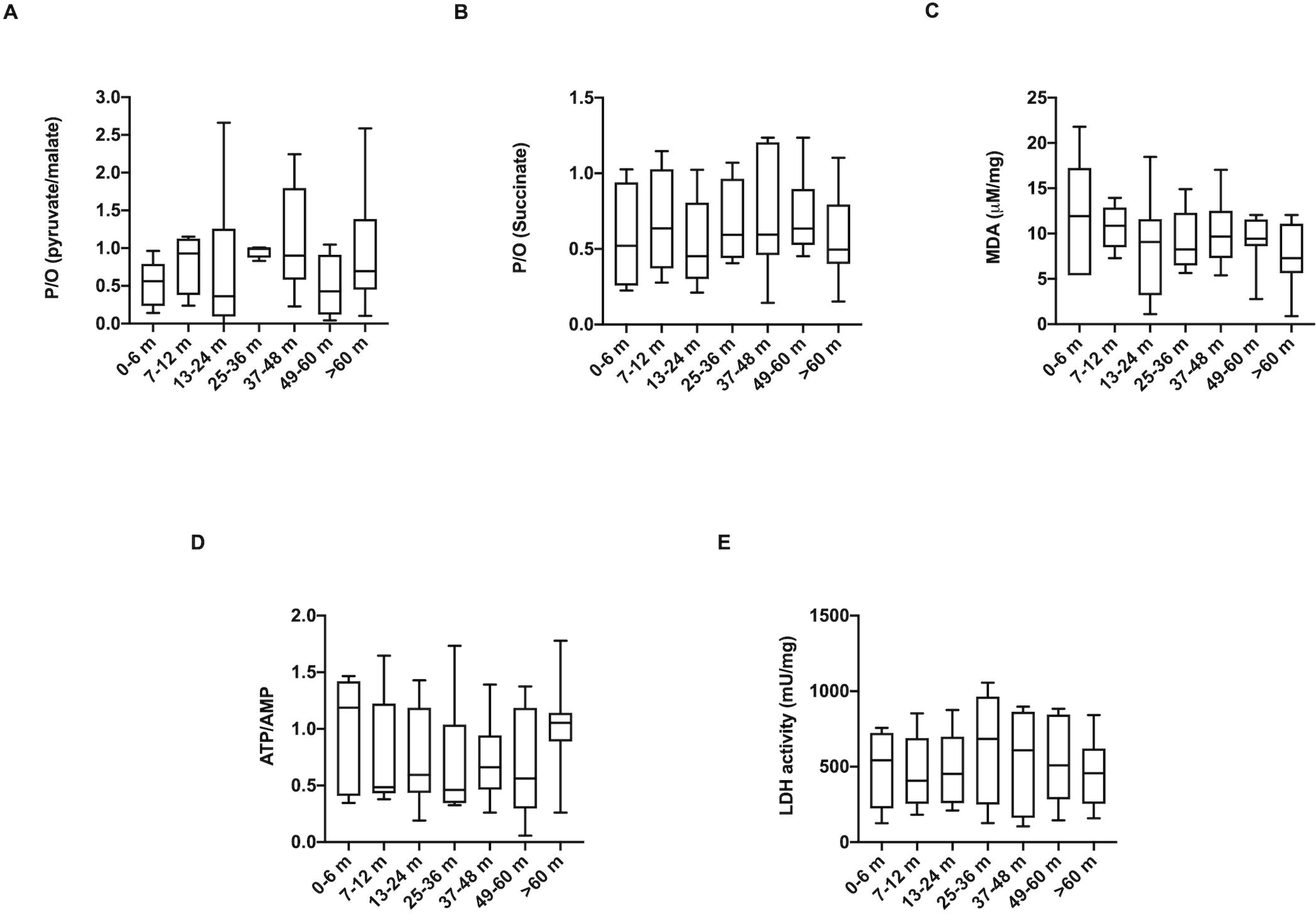
Comparison of glucose metabolism alteration in time after chemo/chemoradiotherapy. In this figure the biochemical parameters evaluated in the MNCs isolated from CCS patients are divided on the basis of the time elapsed since the last clinical treatment at the time of blood collection. The time is indicated in months: 0-6 months (n=12), 7-12 months (n=20), 13-24 months (n=15), 25-36 months (n=34), 37-48 months (n=24), 49-60 months (n=18), >60 months (n=21). Panel A and B reports the P/O value, obtained as the ratio between ATP synthesis and oxygen consumption, in the presence of pyruvate + malate or succinate, respectively. Panel C shows the MDA level, as marker of lipid peroxidation. Panel D reports the cellular energy status, expressed as the ATP/AMP ratio. Panel E shows the LDH activity, as marker of anaerobic glycolysis. No significant differences have been observed among the analyzed samples.

It is important to note that the alterations of glucose metabolism are observed only in the CCS patients undergoing chemo/chemoradiotherapy, while patients with hematological cancer undergoing allogeneic bone marrow transplantation show no metabolic changes compared to the healthy subjects of the same age (Figure 6). From one side this is obvious since the transplanted cells from healthy donors did not experience any chemo/chemoradiotherapy. At the same time this, indirectly, provides important information since it shows that chemo/chemoradiotherapy does not compromise the hematopoietic microenvironment and allows a normal hematopoietic function of incoming transplanted cells.

**Figure 6.**
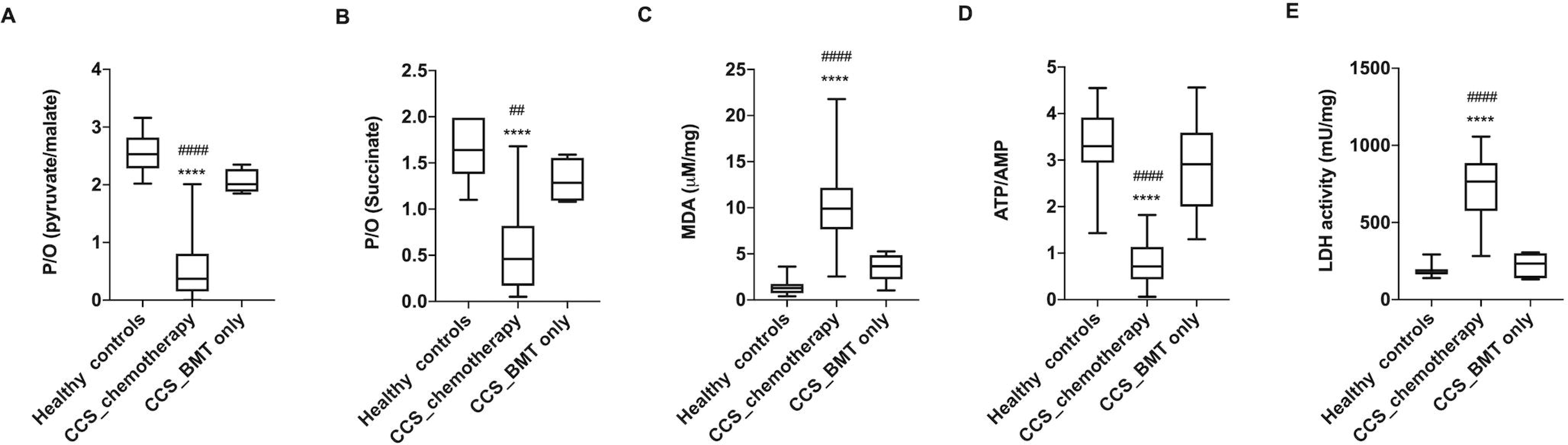
The glucose metabolism alterations are detectable in CCS after chemo/chemoradiotherapy but not after the only bone marrow transplantation. In this figure the reported biochemical parameters are evaluated in the MNCs isolated from hematological CCS patients treated with chemo/chemoradiotherapy (<20 y.o., n=79), CCS patients treated only with bone marrow transplantation (BMT) (<20 y.o., n=15) and the age-matched healthy subjects (n=35) Panel A and B reports the P/O value, obtained as the ratio between ATP synthesis and oxygen consumption, in the presence of pyruvate + malate or succinate, respectively. Panel C shows the MDA level, as marker of lipid peroxidation. Panel D reports the cellular energy status, expressed as the ATP/AMP ratio. Panel E shows the LDH activity, as marker of anaerobic glycolysis. **** indicates significant differences for p<0.0001 between healthy samples and CCS patients treated with chemo/chemoradiotherapy ## and #### indicates significant differences for p<0.0001 between CCS patients treated with only BMT and CCS patients treated with chemo/chemoradiotherapy. No significant differences have been observed between healthy subjects and CCS patients treated with only BMT.

### The metabolic alterations in CCS are suggestive of anticipated aging

Biochemical data suggest that alterations in cellular metabolism, observed in CCS after chemotherapy and radiotherapy, were similar to those observed in elderly subjects. Thus, utilizing our recently developed model to predict age, based on some glucose metabolism parameters (ATP/AMP ratio, P/O ratio in the presence of Pyr+Mal and succinate, MDA level and LDH activity)^28^, we have calculated the age of CCS subjects with hematological and solid tumors and age-matched (< 40 y.o.) or older healthy controls (> 60 y.o.); further we compared those values to the real age of these subjects. As shown in Figure 7 and Supplementary Table 2, the age of healthy subjects computed by the mathematical model is similar to their real age, and the deviation is part of the 9-year model error^28^. In contrast, in CCS samples, the calculated age was significantly higher than the real age.

**Figure 7.**
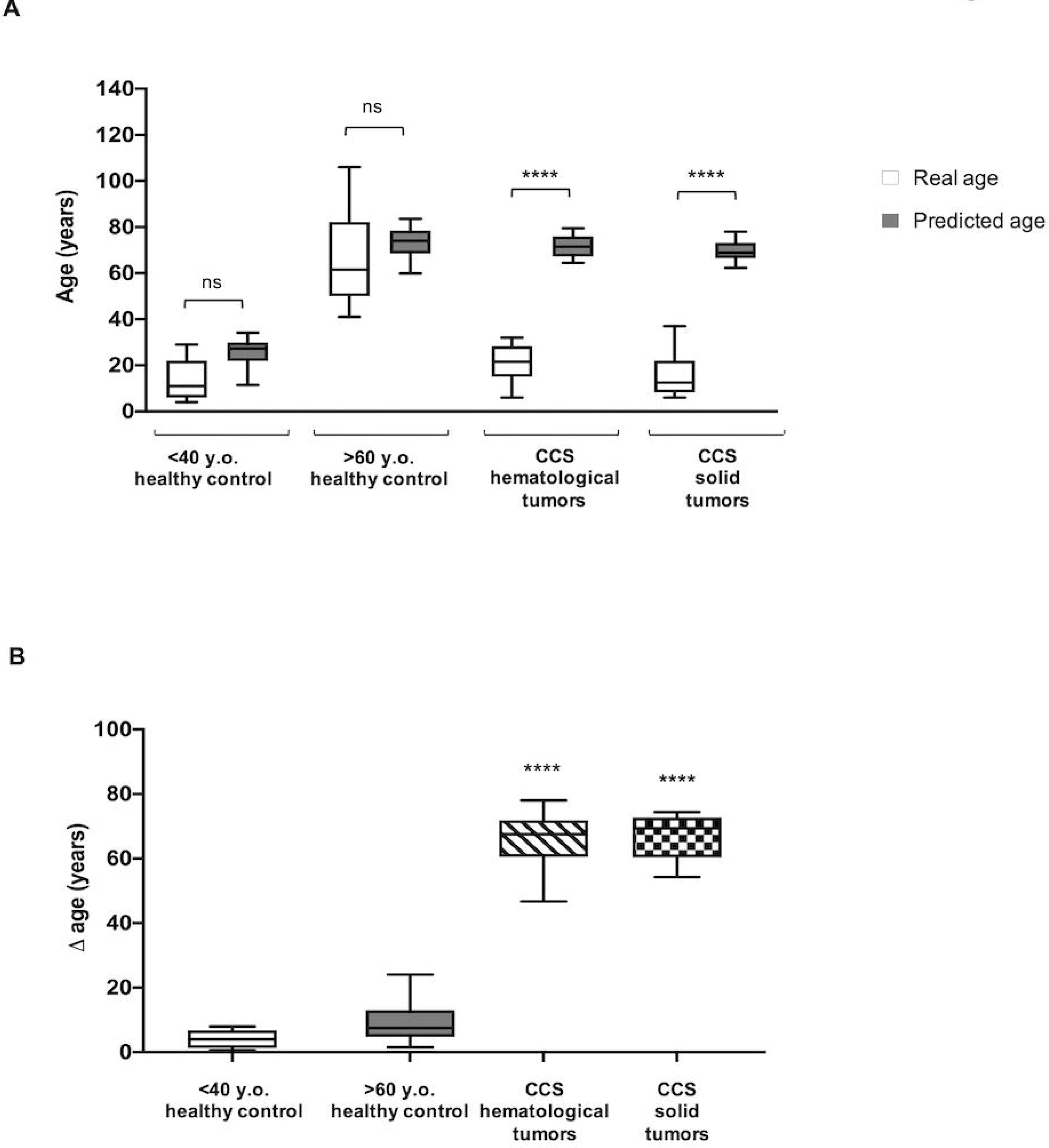
Comparison of real and predicted age in MNC isolated from CCS and in age- matched and elderly healthy control. The predict age was obtained applying a mathematical model developed by machine-learning method and based on several parameters of glucose metabolism^28^. The data are obtained evaluating the biochemical parameters on MNC isolated from: age-matched healthy control (<40 y.o. n= 59), elderly healthy control (>60 y.o. n=47), CCS hematological tumors (<40 y.o. n=89), and CCS solid tumors (<40 y.o. n=55). Panel A show a comparison between the real and predicted age of the same sample. ****: p<0.0001; *ns*: no significant difference between real and predicted age. Panel B reports Δ values calculated between predicted and real age among CCS (hematological and solid tumors), age-matched and elderly controls. ****: p<0.0001; *ns*: no significant differences

### Proteomic analysis confirms alteration of mitochondrial metabolism

The proteomic analysis approach (Figure 8) confirms that CCS samples were characterized by altered glucose metabolism. In particular, data show a higher expression in CCS of some proteins involved in the anaerobic metabolism (LDHB) and in the transport of glucose (SLC2A3 and SLC2A1) than in the age-matched control, confirming the switch from mitochondrial metabolism to anaerobic glycolysis. Reciprocally, the expression of some subunits of respiratory complex I and ATP synthase as well as some enzymes involved in the antioxidant defenses, such as glutathione synthase (GSS), catalase (CAT), and glucose 6-phosphate dehydrogenase (G6PD) appeared downregulated in CCS in comparison to the control.

**Figure 8.**
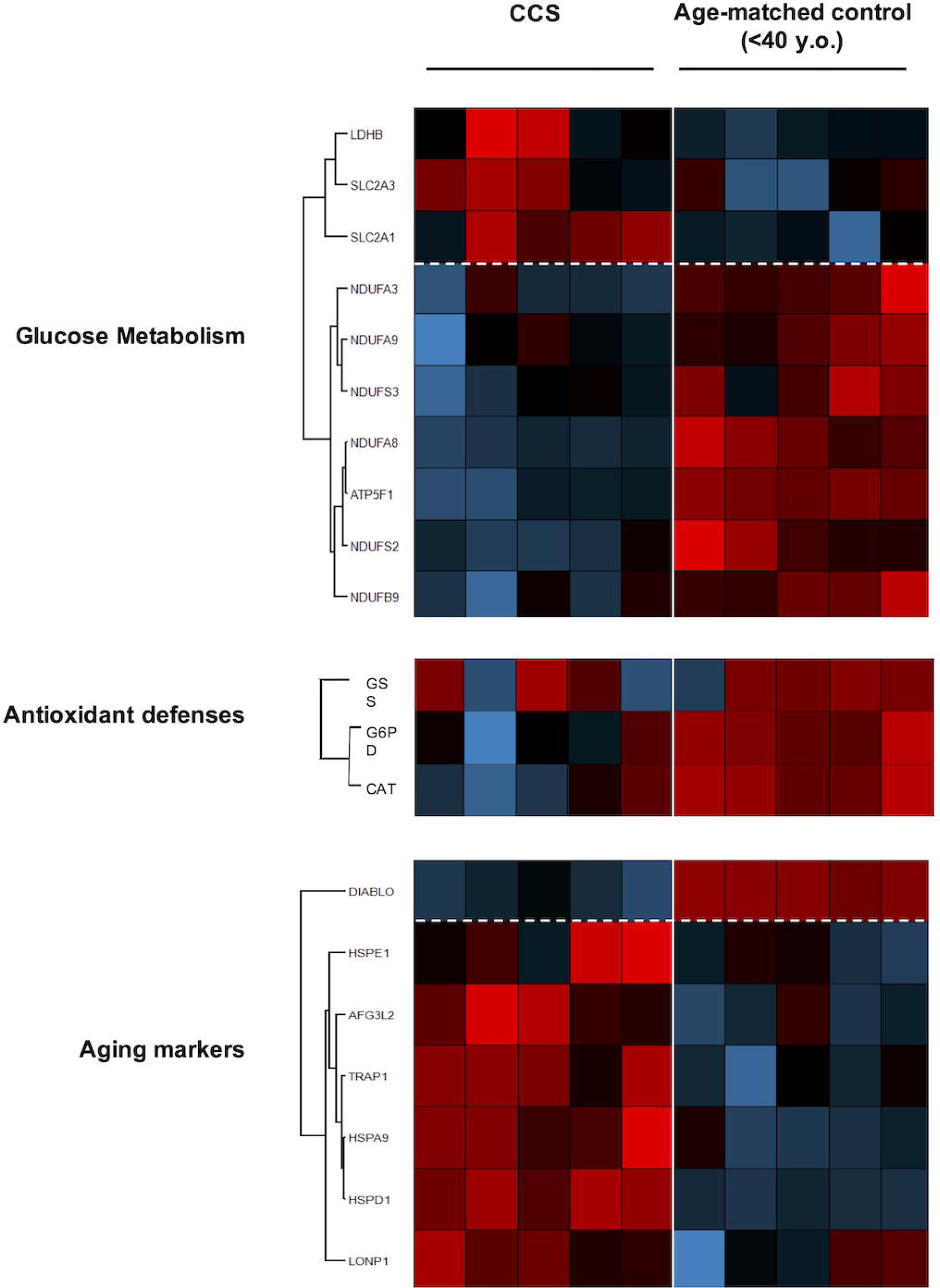
Proteomic analysis on MNC isolated from CCS and age-matched healthy control. The figure shows a representative expression heat maps of proteins involved in glucose metabolism, antioxidant defenses and aging markers. Data are expressed as ratios over mean values for the two conditions (red = expression above mean, black= expression at mean; blue = expression below mean). The data is represented of five independent experiments.

Finally, we investigated the regulation of expression of proteins involved in the aging processes, observing a downregulation of DIABLO, an apoptosis-inducing protein, and the upregulation of proteins involved in stress adaptation (HSPA9), negative modulation of OxPhos (TRAP1), and mitochondrial protein importation, folding and degradation (i.e., HSPD1, HSPE1, AFG3L2, LONP1), which are suggestive of an impaired mitochondrial function.

### Genes involved in mitochondrial biogenesis and activity regulation are altered in CCS

To disclose possible mechanisms involved in the metabolic alterations observed in CCS, the expression of several genes involved in mitochondrial function has been investigated. Our panel includes genes involved in (i) mitochondrial autophagy (PINK1, PARKIN), (ii) mitochondrial fusion/fission processes (MNF1 and FIS1), (iii) mitochondrial biogenesis (CLUH and NRF1), and (iv) regulation of mitochondrial metabolic pathways (PGC1-α, SIRT1and SIRT6). CCS showed significant downregulation of CLUH, PGC1-α, and SIRT6 gene expression, in comparison to any control group (age-matched and elderly), (Figure 9, Panel A). In other words, the altered expression of these genes appears a peculiar sign of CCS, not linked with the “physiological aging”. Conversely, the gene expression of PINK1, PARKIN, MNF1, FIS1, NRF1, and SIRT1 appeared not compromised in CCS and similar to healthy control samples (data not shown).

The qPCR results are confirmed by the WB analysis that shows that the protein expression of CLUH, PGC1-α, and SIRT6in cell lysates were similar in young and elderly healthy subjects but is low detectable in CCS samples (Figure 9, Panel B).

**Figure 9.**
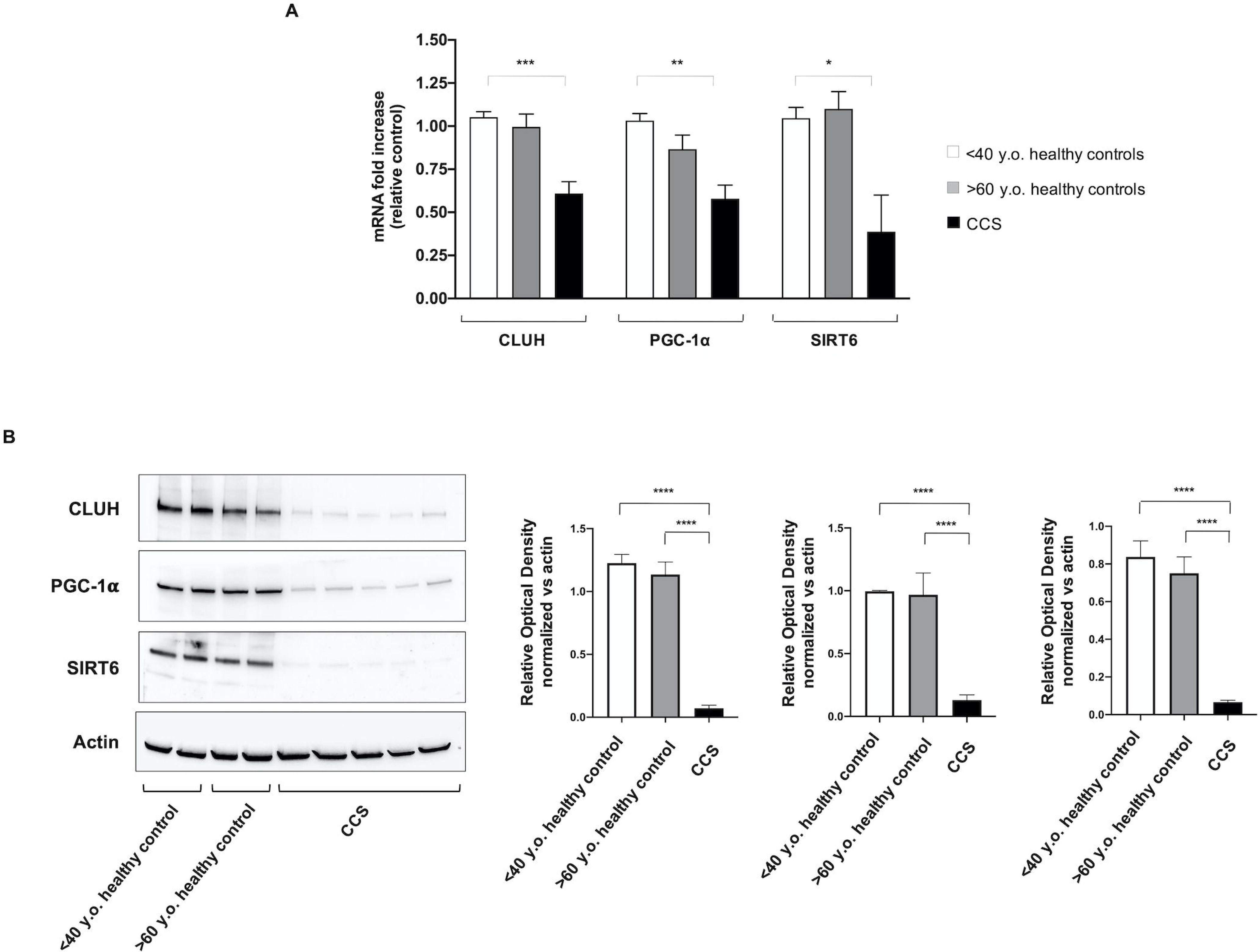
Evaluation of gene and protein expression of CLUH, PGC1-α, and SIRT6 in MNC isolated from CCS and in age-matched and elderly healthy control. In Panel A, the graph reports the genomic expression of CLUH, PGC1-α, and SIRT6 in MNC isolated from age-matched healthy control (<40 y.o. n=10), CCS (<40 y.o. n=16), and elderly healthy control (>60 y.o. n=12). Panel B reports the WB signals against CLUH, PGC1-α, and SIRT6 (left) and the relative densitometric analyses (right) obtained on MNC isolated from age-matched healthy control (<40 y.o.), elderly healthy control (>60 y.o.), and CCS (<40 y.o.). Data are representative of four independent experiments and are expressed as mean ± SD. The values of WB signals are normalized against actin signal. *, **, *** and **** indicate a significant difference for p<0.05, 0.01, 0.001 and 0.0001, respectively, between CCS samples and age-matched healthy controls. No significant differences have been observed between age-matched and elderly healthy controls.

## DISCUSSION

Progress in the treatment of children with cancer has led to a remarkable increase in the number of survivors living into adulthood. Despite this impressive success, the impact on the long term health of these individuals remains substantial, with most experiencing chronic health conditions related to previous treatment ^44^. Late effects of chemo/chemoradiotherapy ^45–47^ are reflected in outcomes typically associated with aging, like minimal cognitive dysfunction, reduced muscle strength, cardiovascular disease, poor exercise tolerance, increased cellular senescence, reduced telomere length, epigenetic modifications, somatic mutations, and mitochondrial damage ^8–14^.

Therefore, the objective of the present work was to identify molecular and biochemical alterations into blood cells in order to give a pathophysiological basis of early aging observed in CCS. Our attention focused on the mitochondrial energy metabolism and the relative oxidative stress production since the reactive oxygen species is one of the principal causes of aging ^48^.

Our data show that CCS-MNCs are characterized by an impairment of OxPhos efficiency, evaluated in term of P/O ratio ad MMP, due to the uncoupled status between the oxygen consumption and ATP synthesis, in comparison to the age-matched control; indeed, the P/O ratio was very similar to those observed in the elderly control (>60 y.o.). This aerobic metabolism inefficiency was observed to be associated with (i) a lower expression of some subunits of Complex I (NDUFA3, NDUFA9, NDUFS3, NDUFA8, NDUFS2, and NDUFB9) and ATP synthase (ATP5F1); (ii) over-expression of several heath-shock proteins (HSP) involved in the mitochondrial protein importation, folding and degradation (i.e. HSPE1 ^49^, HSPD1 ^50^, AFG3L2 ^51^, LONP1 ^52^), as well as in stress adaptation (i.e. HSPA9 ^53^).

The metabolic switch observed in our analysis is further confirmed by the increased expression of TRAP1, a negative modulator of OxPhos metabolism, and an inducer of glycolysis ^54^. By contrast, the expression of DIABLO, a protein favoring the cellular apoptosis ^55^, appears lower with respect to the age-matched control cells, confirming that the programmed cell death is impaired in senescent cells ^56^. Thus, the accelerated aging in CCS could be associated with the accumulation of senescent cells ^57^, which may explain the frailty observed in CCS, since the accumulation of senescent cells for prolonged periods ^57^ favor the onset of age-related diseases because of their low, but chronic, senescence-associated secretory phenotype ^57–59^.

The CCS-MNC metabolic alteration determines the reduction of chemical energy synthesis, influencing the ATP/AMP ratio. Moreover, mitochondrial inefficiency increases oxidative stress production, as confirmed by the high ROS level and the evident accumulation of MDA, a final product of lipid peroxidation, suggesting that the cellular antioxidant defenses are not sufficient to counteract the increased oxidative stress. This hypothesis is further corroborated by the proteomic analysis that shows a lower expression of G6PD, GSS, and CAT, enzymes involved in the antioxidant response. Moreover, considering that the integrity of the inner mitochondrial membrane is essential for an efficient OxPhos and that mitochondria are major sources of oxidative stress, the decrement of antioxidant defenses associated with the localized oxidative stress production could cause structural damages in the inner mitochondrial membrane, exacerbating the ROS production. In turn, this could trigger a vicious circle in which ROS production becomes both cause and effect of mitochondrial dysfunction ^60^. Of note, the OxPhos inefficiency is more evident in the presence of pyruvate/malate than with the induction by succinate. This could be explained considering that pyruvate + malate activates the pathway triggered by Complexes I that is the principal source of ROS production ^61, 62^. Conversely, succinate, stimulating Complex II, is less involved in oxidative stress production.

In response to the mitochondrial dysfunction, CCS-MNCs increase the expression and the activity of LDH and the expression of the glucose carriers (SLC2A3 and SLC2A1), to compensate the altered energy balance. This switch from aerobic to anaerobic metabolism is imposed by two cellular needs: (i) the restoration, at least in part, of the ATP levels, by incrementing the glucose catabolism; (ii) the balancing of the NADH/NAD^+^ ratio. The recycling of NADH to NAD^+^ is essential to avoid glycolysis slow-down ^63^, and reductive stress accumulation, which could increment oxidative stress ^64^. In fact, in healthy cells, the correct NADH/NAD^+^ ratio is mainly maintained by the respiratory complex I activity, but in the case of mitochondrial dysfunction, this task is performed exclusively by LDH ^63^.

To further extend the investigation on the possible causes of the metabolic mitochondria alteration, a panel of genes involved in mitochondrial biogenesis, function and regulation have been evaluated. Results show that the genes involved in mitophagy, fusion and fission processes (PINK1, PARKIN, MFN1, FIS1, and NRF1) seem to be not impaired with respect to the controls, while genes involved in mitochondrial function, such as CLUH, PCG1-, and SIRT6, appear remarkably less expressed. These data are confirmed by the analysis of protein expression. SIRT6 and PGC1-α play a pivotal role in the regulation of energy metabolism: SIRT6 determines a low level of HIF-α, favoring the aerobic metabolism^65^; whilePGC1-α modulates both the biogenesis and the composition and functions of individual mitochondria^66^. CLUH is a gene codifying for a cytosolic RNA-binding protein involved in the distribution of mitochondria inside the cell and linked to the efficiency of mitochondria OxPhos ^32, 67^. Therefore, the CLUH low expression is in line with the poor OxPhos activity observed by biochemical analysis. Moreover, these results corroborate previous observations on the association between CLUH and mitochondria function ^32, 67, 68^. Interestingly, the alterations of CLUH, SIRT6, and PGC1-α are not observed in elderly controls, despite the metabolic impairment, suggesting that the biochemical mechanism of aging runs in different ways, at least in some relevant aspects, in CCS compared to the aging as a “physiological process”.

The early aging of CCS has been confirmed by the application of our mathematical model based on glucose metabolism and oxidative stress parameters ^28^. In particular, results show that the predicted age is very different in comparison to the real age in CCS patients treated both for hematological and solid tumors. In contrast, by utilizing our mathematical model, the predicted age of healthy controls seems not different to the real age.

Making a general consideration of the data reported herein, it is interesting to note that metabolic dysfunctions and genetic alterations are irrespective of heterogeneity of the original disease, the relative treatment received, and the time elapsed between the last therapy and the time of biochemical and genetic analysis. In other words, the cellular biochemical/molecular lesions observed among the 196 CCS examined was consistent and rather homogeneous, not allowing us to correlate the presence or the severity of frailty/anticipated aging symptoms with these biochemical markers. However, although the analysis of MNC represents a good model to evaluate the health status of the entire organism ^29^, it is important to consider that the lifespan and the mitochondrial density of MNCs is different in comparison to those of other organs, such as liver and heart ^69^. This could imply that other tissues could accumulate a different degree of damage from the alteration of the mitochondrial metabolism. It is possible that other tissues may harbor biochemical/molecular alterations of different severity depending on the combination of tissue specificity with the type of therapy and the time elapsed between it and the analysis of biochemical parameters. However, in this work we have not evaluated cells of tissues other than MNC that remains, by far, the most accessible cell source.

In our study, there is no evidence of improvement of the biochemical injury with time. In fact, the same damage is observed in CCS examined few years after the termination of therapy and in patients who had stopped their treatments more than 10 years before being enrolled in this study. This suggests that there is permanent damage in hematopoietic stem cells (HSC) and, apparently, there are not spared HSC that can replace the damaged ones with time. Unfortunately, because the riddle of whether HSC are all active at the same time has not yet been solved ^70, 71^ it is very difficult to address this topic.

A general better response to chemotherapy and a higher treatment-related toxicity are both observed in childhood cancer patients than in adults. This could be, at least in part, ascribed to a higher expression of the apoptotic protein machinery in infant tissues than in adulthood ^72^. Nevertheless, this holds true for liver, brain, heart and kidney tissues, but not for MNC cells ^72^. The latter display indeed a high mitochondrial priming for apoptotic death in adults as well. Accordingly, we may hypothesize that extending this study to the adult cancer survivor counterpart might likely disclose long-term treatment-related mitochondrial dysfunction and cellular aging of MNC.

In conclusion, our data suggest that one of the causes of early aging in CCS could be the alteration of aerobic mitochondrial metabolism could be related to a defect in the organization of their biogenesis and metabolism modulation. Moreover, the evaluation of glucose metabolism may represent a new no-invasive tool to detect precociously the aging of CCS related to chemo/radiotherapy. These initial findings warrant further research aimed to discover additional molecular/biochemical alterations enabling to disclose more severe from less severe conditions at cellular level and possibly correlating them with clinical symptoms.

Finally, this study may facilitate the possibility to identify therapies to restore the mitochondrial function, slow downing the aging and the associated pathological conditions in CCS.

## REFERENCES

1. Bhatia S, Meadows AT: Long-term follow-up of childhood cancer survivors: future directions for clinical care and research. Pediatr Blood Cancer 46:143–8, 2006

2. Gatta G, Capocaccia R, Stiller C, et al: Childhood cancer survival trends in Europe: a EUROCARE Working Group study. J Clin Oncol 23:3742–51, 2005

3. Shapiro CL: Cancer Survivorship. N Engl J Med 379:2438–2450, 2018

4. Nelson MB, Meeske K: Recognizing health risks in childhood cancer survivors. J Am Acad Nurse Pract 17:96–103, 2005

5. Landier W, Skinner R, Wallace WH, et al: Surveillance for Late Effects in Childhood Cancer Survivors. J Clin Oncol 36:2216–2222, 2018

6. Grabow D, Kaiser M, Hjorth L, et al: The PanCareSurFup cohort of 83,333 five-year survivors of childhood cancer: a cohort from 12 European countries. Eur J Epidemiol 33:335–349, 2018

7. Friedman DL, Whitton J, Leisenring W, et al: Subsequent neoplasms in 5-year survivors of childhood cancer: the Childhood Cancer Survivor Study. J Natl Cancer Inst 102:1083–95, 2010

8. Henderson TO, Ness KK, Cohen HJ: Accelerated aging among cancer survivors: from pediatrics to geriatrics. Am Soc Clin Oncol Educ Book e423–30, 2014

9. Ness KK, Krull KR, Jones KE, et al: Physiologic frailty as a sign of accelerated aging among adult survivors of childhood cancer: a report from the St Jude Lifetime cohort study. J Clin Oncol 31:4496–503, 2013

10. Armstrong GT, Kawashima T, Leisenring W, et al: Aging and risk of severe, disabling, life-threatening, and fatal events in the childhood cancer survivor study. J Clin Oncol 32:1218–27, 2014

11. Hartman A, van den Bos C, Stijnen T, et al: Decrease in peripheral muscle strength and ankle dorsiflexion as long-term side effects of treatment for childhood cancer. Pediatr Blood Cancer 50:833–7, 2008

12. Ness KK, Baker KS, Dengel DR, et al: Body composition, muscle strength deficits and mobility limitations in adult survivors of childhood acute lymphoblastic leukemia. Pediatr Blood Cancer 49:975–81, 2007

13. Krull KR, Sabin ND, Reddick WE, et al: Neurocognitive function and CNS integrity in adult survivors of childhood hodgkin lymphoma. J Clin Oncol 30:3618–24, 2012

14. Schuitema I, Deprez S, Van Hecke W, et al: Accelerated aging, decreased white matter integrity, and associated neuropsychological dysfunction 25 years after pediatric lymphoid malignancies. J Clin Oncol 31:3378–88, 2013

15. Ness KK, Armstrong GT, Kundu M, et al: Frailty in childhood cancer survivors. Cancer 121:1540–1547, 2015

16. Kadan-Lottick NS, Zeltzer LK, Liu Q, et al: Neurocognitive functioning in adult survivors of childhood non-central nervous system cancers. J Natl Cancer Inst 102:881–893, 2010

17. Meeske KA, Nelson MB: The role of the long-term follow-up clinic in discovering new emerging late effects in adult survivors of childhood cancer. J Pediatr Oncol Nurs 25:213–9, 2008

18. Lipshultz SE, Franco VI, Miller TL, et al: Cardiovascular Disease in Adult Survivors of Childhood Cancer. Annu Rev Med 66:161–176, 2015

19. Armstrong GT, Liu Q, Yasui Y, et al: Late Mortality Among 5-Year Survivors of Childhood Cancer: A Summary From the Childhood Cancer Survivor Study. J Clin Oncol 27:2328–2338, 2009

20. Ness KK, Kirkland JL, Gramatges MM, et al: Premature Physiologic Aging as a Paradigm for Understanding Increased Risk of Adverse Health Across the Lifespan of Survivors of Childhood Cancer. J Clin Oncol 36:2206–2215, 2018

21. Lipshultz SE, Franco VI, Miller TL, et al: Cardiovascular Disease in Adult Survivors of Childhood Cancer. Annu Rev Med 66:161–176, 2015

22. Horvath S: DNA methylation age of human tissues and cell types. Genome Biol 14:R115, 2013

23. López-Otín C, Blasco MA, Partridge L, et al: The hallmarks of aging. Cell 153:1194–217, 2013

24. Sastre, Federico V. Pallardo, Jos & J, Pallardó F V, Viña J: Mitochondrial Oxidative Stress Plays a Key Role in Aging and Apoptosis. IUBMB Life (International Union Biochem Mol Biol Life) 49:427–435, 2000

25. Correia□ Melo C, Marques FD, Anderson R, et al: Mitochondria are required for pro□ageing features of the senescent phenotype. EMBO J 35:724–742, 2016

26. Fakouri NB, Hou Y, Demarest TG, et al: Toward understanding genomic instability, mitochondrial dysfunction and aging. FEBS J 286:1058–1073, 2019

27. Korolchuk VI, Miwa S, Carroll B, et al: Mitochondria in Cell Senescence: Is Mitophagy the Weakest Link? EBioMedicine 21:7–13, 2017

28. Ravera S, Podestà M, Sabatini F, et al: Discrete Changes in Glucose Metabolism Define Aging. Sci Rep 9:10347, 2019

29. McKerrell T, Park N, Moreno T, et al: Leukemia-Associated Somatic Mutations Drive Distinct Patterns of Age-Related Clonal Hemopoiesis. Cell Rep 10:1239–1245, 2015

30. Cappelli E, Cuccarolo P, Stroppiana G, et al: Defects in mitochondrial energetic function compels Fanconi Anaemia cells to glycolytic metabolism. Biochim Biophys Acta - Mol Basis Dis 1863:1214–1221, 2017

31. Hinkle PC: P/O ratios of mitochondrial oxidative phosphorylation. Biochim Biophys Acta 1706:1–11, 2005

32. Ravera S, Podestà M, Sabatini F, et al: Mesenchymal stem cells from preterm to term newborns undergo a significant switch from anaerobic glycolysis to the oxidative phosphorylation. Cell Mol Life Sci 75:889–903, 2018

33. Ravera S, Dufour C, Cesaro S, et al: Evaluation of energy metabolism and calcium homeostasis in cells affected by Shwachman-Diamond syndrome. Sci Rep 6:25441, 2016

34. Bruno S, Ghiotto F, Tenca C, et al: N-(4-hydroxyphenyl)retinamide promotes apoptosis of resting and proliferating B-cell chronic lymphocytic leukemia cells and potentiates fludarabine and ABT-737 cytotoxicity. Leukemia 26:2260–2268, 2012

35. Cappelli E, Degan P, Bruno S, et al: The passage from bone marrow niche to bloodstream triggers the metabolic impairment in Fanconi Anemia mononuclear cells. Redox Biol 36:101618, 2020

36. Kulak NA, Pichler G, Paron I, et al: Minimal, encapsulated proteomic-sample processing applied to copy-number estimation in eukaryotic cells. Nat Methods 11:319–24, 2014

37. Tyanova S, Temu T, Cox J: The MaxQuant computational platform for mass spectrometry-based shotgun proteomics. Nat Protoc 11:2301–2319, 2016

38. Tyanova S, Temu T, Sinitcyn P, et al: The Perseus computational platform for comprehensive analysis of (prote)omics data. Nat Methods 13:731–40, 2016

39. Vizcaíno JA, Csordas A, del-Toro N, et al: 2016 update of the PRIDE database and its related tools. Nucleic Acids Res 44:D447–56, 2016

40. Bradford MM: A rapid and sensitive method for the quantitation of microgram quantities of protein utilizing the principle of protein-dye binding. Anal Biochem 72:248–254, 1976

41. Tiwari BS, Belenghi B, Levine A: Oxidative stress increased respiration and generation of reactive oxygen species, resulting in ATP depletion, opening of mitochondrial permeability transition, and programmed cell death. Plant Physiol 128:1271–81, 2002

42. Cui H, Kong Y, Zhang H: Oxidative stress, mitochondrial dysfunction, and aging. J Signal Transduct 2012:646354, 2012

43. Srivastava S: The Mitochondrial Basis of Aging and Age-Related Disorders. Genes (Basel) 8:398, 2017

44. Bhakta N, Liu Q, Ness KK, et al: The cumulative burden of surviving childhood cancer: an initial report from the St Jude Lifetime Cohort Study (SJLIFE). Lancet (London, England) 390:2569–2582, 2017

45. Vassal G, Schrappe M, Pritchard-Jones K, et al: The SIOPE strategic plan: A European cancer plan for children and adolescents. J Cancer Policy 8:17–32, 2016

46. Oeffinger KC, Mertens AC, Sklar CA, et al: Chronic Health Conditions in Adult Survivors of Childhood Cancer. N Engl J Med 355:1572–1582, 2006

47. Geenen MM, Cardous-Ubbink MC, Kremer LCM, et al: Medical Assessment of Adverse Health Outcomes in Long-term Survivors of Childhood Cancer. JAMA 297:2705, 2007

48. Dai D-F, Chiao YA, Marcinek DJ, et al: Mitochondrial oxidative stress in aging and healthspan. Longev Heal 3:6, 2014

49. Bross P, Li Z, Hansen J, et al: Single-nucleotide variations in the genes encoding the mitochondrial Hsp60/Hsp10 chaperone system and their disease-causing potential. J Hum Genet 52:56–65, 2007

50. Bross P, Fernandez-Guerra P: Disease-Associated Mutations in the HSPD1 Gene Encoding the Large Subunit of the Mitochondrial HSP60/HSP10 Chaperonin Complex. Front Mol Biosci 3:49, 2016

51. Jadiya P, Tomar D: Mitochondrial Protein Quality Control Mechanisms. Genes (Basel) 11, 2020

52. Bezawork-Geleta A, Brodie EJ, Dougan DA, et al: LON is the master protease that protects against protein aggregation in human mitochondria through direct degradation of misfolded proteins. Sci Rep 5:17397, 2015

53. Starenki D, Sosonkina N, Hong S-K, et al: Mortalin (GRP75/HSPA9) Promotes Survival and Proliferation of Thyroid Carcinoma Cells. Int J Mol Sci 20, 2019

54. Yoshida S, Tsutsumi S, Muhlebach G, et al: Molecular chaperone TRAP1 regulates a metabolic switch between mitochondrial respiration and aerobic glycolysis. Proc Natl Acad Sci U S A 110:E1604–12, 2013

55. Redza-Dutordoir M, Averill-Bates DA: Activation of apoptosis signalling pathways by reactive oxygen species. Biochim Biophys Acta - Mol Cell Res 1863:2977–2992, 2016

56. Wang E: Senescent human fibroblasts resist programmed cell death, and failure to suppress bcl2 is involved. Cancer Res 55:2284–92, 1995

57. Krishnamurthy J, Torrice C, Ramsey MR, et al: Ink4a/Arf expression is a biomarker of aging. J Clin Invest 114:1299–307, 2004

58. Baar MP, Brandt RMC, Putavet DA, et al: Targeted Apoptosis of Senescent Cells Restores Tissue Homeostasis in Response to Chemotoxicity and Aging. Cell 169:132–147.e16, 2017

59. de Keizer PLJ: The Fountain of Youth by Targeting Senescent Cells? Trends Mol Med 23:6–17, 2017

60. Sanz A, Caro P, Gómez J, et al: Testing the vicious cycle theory of mitochondrial ROS production: effects of H2O2 and cumene hydroperoxide treatment on heart mitochondria. J Bioenerg Biomembr 38:121–127, 2006

61. Kausar S, Wang F, Cui H: The Role of Mitochondria in Reactive Oxygen Species Generation and Its Implications for Neurodegenerative Diseases. Cells 7:274, 2018

62. Hirst J, King MS, Pryde KR: The production of reactive oxygen species by complex I, in Biochemical Society Transactions. 2008, pp 976–980

63. Cantó C, Menzies KJ, Auwerx J: NAD(+) Metabolism and the Control of Energy Homeostasis: A Balancing Act between Mitochondria and the Nucleus. Cell Metab 22:31–53, 2015

64. Korge P, Calmettes G, Weiss JN: Increased reactive oxygen species production during reductive stress: The roles of mitochondrial glutathione and thioredoxin reductases. Biochim Biophys Acta - Bioenerg 1847:514–525, 2015

65. Sociali G, Magnone M, Ravera S, et al: Pharmacological Sirt6 inhibition improves glucose tolerance in a type 2 diabetes mouse model. FASEB J 31, 2017

66. Austin S, St-Pierre J: PGC1a and mitochondrial metabolism-emerging concepts and relevance in ageing and neurodegenerative disorders. J Cell Sci 125:4963–4971

67. Gao J, Schatton D, Martinelli P, et al: CLUH regulates mitochondrial biogenesis by binding mRNAs of nuclear-encoded mitochondrial proteins. J Cell Biol 207:213–23, 2014

68. Schatton D, Pla-Martin D, Marx M-C, et al: CLUH regulates mitochondrial metabolism by controlling translation and decay of target mRNAs. J Cell Biol 216:675–693, 2017

69. Veltri KL, Espiritu M, Singh G: Distinct genomic copy number in mitochondria of different mammalian organs. J Cell Physiol 143:160–164, 1990

70. Qiu J, Papatsenko D, Niu X, et al: Divisional history and hematopoietic stem cell function during homeostasis. Stem cell reports 2:473–90, 2014

71. Lu R: Sleeping Beauty Wakes Up the Clonal Succession Model for Homeostatic Hematopoiesis. Cell Stem Cell 15:677–678, 2014

72. Sarosiek KA, Fraser C, Muthalagu N, et al: Developmental Regulation of Mitochondrial Apoptosis by c-Myc Governs Age- and Tissue-Specific Sensitivity to Cancer Therapeutics. Cancer Cell 31:142–156, 2017

