## Supplemental Table 1 for "Identification of Biochemical and Molecular Markers of Early Aging in Childhood Cancer Survivors"

**Table 1_Supplementary – Metabolism of CCS patients and controls**

|  | | **age**  **<10 years** | **age**  **11-20 years** | **age**  **21-40 years** | **age**  **<10 years** | **age**  **11-20 years** | **Age**  **21-40 years** | **age**  **<10 years** | **age**  **11-20 years** | **age**  **21-40 years** | **age**  **41-60 years** | **age**  **61-80 years** | **age**  **>80 years** |
| --- | --- | --- | --- | --- | --- | --- | --- | --- | --- | --- | --- | --- | --- |
|  |  | **CCS patients**  **(hematological tumors)** | | | **CCS patients**  **(solid tumors)** | | | **Age-matched**  **healthy donors** | | | **Older**  **healthy donors** | | |
| **P/O**  **Pyruvate/Malate** | **Range** | 0-4.2 | 0-3.09 | 0.37-2.59 | 0.44-3.66 | 0.26-1.83 | 0.25-1.77 | 2.04-3.16 | 1.09-3.08 | 2.11-2.85 | 0-3.11 | 0-1.42 | 0-1.9 |
|  | **median±SD** | 0.64±0.29 | 0.6±0.09 | 1.08±0.23 | 0.79±0.24 | 0.95±0.14 | 0.53±0.3 | 2.53±0.13 | 2.51±0.16 | 2.4±0.07 | 1.67±0.17 | 0.61±0.09 | 0.32±0.08 |
| **P/O**  **Succinate** | **Range** | 0-1.59 | 0-1.45 | 0.63-1.68 | 0.37-1.17 | 0.15-1.29 | 0.14-1.86 | 1.1-1.99 | 0.73-1.77 | 0.8-1.94 | 0.17-1.73 | 0.19-1.42 | 0.24-3.24 |
|  | **median±SD** | 0.23±0.12 | 0.47±0.06 | 0.69±0.19 | 0,6±0.07 | 0.47±0.13 | 0.29±0.40 | 1.64±0.12 | 1.43±0.08 | 1.38±0.13 | 1.43±0.13 | 0.53±0.09 | 0.43±0.13 |
| **ATP/AMP** | **Range** | 0.19-3.26 | 0.06-4 | 0.26-3.08 | 0.2-5.5 | 0.38-3.8 | 0.28-1.6 | 2.9-4.4 | 1.43-4.55 | 2.06-4.4 | 0.61-4.3 | 0.18-2.63 | 0.37-2.08 |
|  | **median±SD** | 0.93±0.14 | 0.84±0.08 | 0.67±0.20 | 0.80±0.18 | 0.99±0.17 | 0.92±0.16 | 3.67±0.17 | 3±0.26 | 2.8±0.27 | 1.12±0.18 | 0.99±0.12 | 0.92±0.08 |
| **LDH (U/mg)** | **Range** | 24.9-885.9 | 8-1056 | 131.8-850.5 | 281.4-964.2 | 246-955 | 762.8-975.3 | 144.7-202.6 | 139.9-292.6 | 102.9-305.5 | 112.5-326.5 | 245.6-371.3 | 294.2-426.3 |
|  | **median±SD** | 426±70.2 | 605.2±43.2 | 450.8±60.7 | 758.8±678 | 598.4±64.2 | 869.1±61.3 | 192.9±7.4 | 181.7±13.7 | 162.3±25.1 | 263.5±11.1 | 291.4±7.9 | 385.1±9.4 |
| **MDA (μM/mg)** | **Range** | 2.07-17.04 | 0.88-24.41 | 3.49-15.14 | 1.12-17.75 | 2.31-28.68 | 2.31-14.66 | 0.62-1.96 | 0.18-2.94 | 0.62-3.17 | 0.85-16.33 | 5.97-12.75 | 8.07-13.20 |
|  | **median±SD** | 8.96±0.83 | 9.67±0.53 | 8.01±0.80 | 7.77±0.72 | 8.60±1.07 | 9.43±1.57 | 1.07±0.19 | 1.29±0.26 | 1.19±0.40 | 5.84±1.33 | 9.74±0.51 | 11.53±0.29 |
