## Supplemental Table 2 for "Identification of Biochemical and Molecular Markers of Early Aging in Childhood Cancer Survivors"

**Table 2 Supplementary – Predicted age *vs* real age in CCS patients and controls**

|  | | **CCS patients**  **Hematological tumors** | **CCS patients**  **Solid tumors** | **Age-matched**  **healthy donors** | **Older**  **healthy donors** |
| --- | --- | --- | --- | --- | --- |
| **Predicted age** | **Range** | 64,47-79,49 | 62,3-77,95 | 15,48-44,11 | 15,62-83,58 |
|  | **median±SE** | 71,92±0,71 | 70,25±1,35 | 23,75±1,14 | 72,4±2,49 |
| **Real age** | **Range** | 4-23 | 6-24 | 8-29 | 31-106 |
|  | **median±SE** | 14,5±1 | 9,5±1,52 | 13±1,38 | 72,5±2,99 |
| **Δ age** | **Range** | 41,47-71,39 | 50,28-67,2 | 0,55-33,11 | 0,1-36,38 |
|  | **median±SE** | 59,06±1,43 | 61,07±1,73 | 6,17±1,33 | 9,06±1,42 |
